# Local regulation of extracellular vesicle traffic by the synaptic endocytic machinery

**DOI:** 10.1101/2021.08.04.454987

**Authors:** Cassandra R. Blanchette, Amy L. Scalera, Kathryn P. Harris, Zechuan Zhao, Kate Koles, Anna Yeh, Julia K. Apiki, Bryan A. Stewart, Avital A. Rodal

**Affiliations:** Department of Biology, Brandeis University, Waltham, MA; Department of Biology, University of Toronto Mississauga, Mississauga, Canada; Department of Cell and Systems Biology University of Toronto, Toronto, Canada

**Keywords:** Amyloid Precursor Protein, Synaptotagmin-4, retromer, Dynactin, Rab11, *Drosophila*, extracellular vesicle, endosome, exosome, Nwk, FCHSD2, clathrin, AP-2, endocytosis, synapse

## Abstract

Neuronal extracellular vesicles (EVs) can be locally released from presynaptic terminals, carrying cargoes that are important in intercellular signaling and disease. EVs are derived from endosomes, but it remains unclear how synaptic cargoes are directed to the EV pathway, rather than undergoing conventional retrograde endosomal transport and degradation. Here, we find that the clathrin-mediated endocytic machinery plays an unexpected role in maintaining a release-competent pool of synaptic EV cargoes. Endocytic mutants, including *nervous wreck* (*nwk*), *Shibire*/Dynamin, and *AP-2*, exhibit local depletion of multiple cargoes in EV donor terminals. Accordingly, *nwk* mutants phenocopy synaptic plasticity defects associated with loss of the EV cargo Synaptotagmin-4, and suppress lethality upon overexpression of the EV cargo Amyloid Precursor Protein. These EV defects are genetically separable from canonical functions of endocytic proteins in synaptic vesicle recycling and synaptic growth. This endocytic pathway opposes the endosomal retromer complex to regulate EV cargo levels, and acts upstream of synaptic cargo removal by retrograde axonal transport. Our data suggest a novel molecular mechanism that protects EV cargoes from local depletion at synapses.

## Introduction

Neurons depend on complex and highly interconnected endosomal membrane trafficking pathways to sort physiologically and pathologically relevant cargoes (Winckler et al., 2018; Yarwood et al., 2020). One function of endosomal trafficking in neurons is to sort cargoes for release in extracellular vesicles (EVs) (Blanchette and Rodal, 2020). EVs are small membrane-bound compartments that transport protein, lipid, and nucleic acid cargoes from EV-releasing cells to target cells (van Niel et al., 2018). In the nervous system, EV-mediated cargo transport regulates intercellular communication and contributes to neurodegenerative disease pathology (Budnik et al., 2016; Holm et al., 2018; Song et al., 2020). However, our understanding of how EV cargo traffic is spatially and temporally regulated within the polarized and complex morphology of neurons remains limited (Blanchette and Rodal, 2020).

Exosomes are a type of EV generated when endosomal multivesicular bodies (MVBs) undergo fusion with the plasma membrane to release their intralumenal vesicles. Alternatively, MVBs can be trafficked to the lysosome for cargo degradation (van Niel et al., 2018). It remains unclear how and where MVBs are generated in neurons, and how they are directed to an EV versus lysosomal fate. We previously described a trafficking pathway that controls levels of EV cargoes in donor neuron synaptic terminals, regulated by endosome-plasma membrane recycling machinery including the retromer complex and Rab11 (Walsh et al., 2021). This suggests the existence of a specialized presynaptic “EV-permissive” recycling compartment, where cargo is protected from degradation *en route* to MVB formation and release from donor cells. However, the mechanisms by which this compartment is loaded are unknown.

The *Drosophila* larval neuromuscular junction (NMJ) is a powerful *in vivo* model for addressing gaps in our understanding of synaptic EV biology (Ashley et al., 2018; Koles et al., 2012; Korkut et al., 2009; Korkut et al., 2013; Walsh et al., 2021). EVs are released from the presynaptic terminals of larval motor neurons and accumulate postsynaptically within the folds of the muscle membrane subsynaptic reticulum (SSR) or are taken up by muscles and glia (Fuentes-Medel et al., 2009; Koles et al., 2012). EVs released at the *Drosophila* NMJ likely represent exosomes, as presynaptic terminals contain MVBs positive for EV cargoes, and release EVs similar in size to exosomes (Koles et al., 2012; Korkut et al., 2009; Walsh et al., 2021). Several endogenously and exogenously expressed transmembrane proteins have been characterized as neuronal EV cargoes at the *Drosophila* NMJ, including Synaptotagmin-4 (Syt4), which drives functional and structural plasticity, as well as the Alzheimer’s disease-associated human Amyloid Precursor Protein (hAPP) (Korkut et al., 2013; Walsh et al., 2021). Thus, the *Drosophila* NMJ provides a unique opportunity to interrogate the mechanisms and functional significance of traffic of EV cargoes in an intact nervous system. Here, using this system, we have identified a novel and unexpected role for clathrin-mediated endocytic machinery in locally maintaining EV cargoes in EV-permissive compartments at synaptic terminals, therefore promoting their release in synaptically-derived EVs. Our work suggests that endocytic machinery regulates endosomal sorting in neurons to protect EV cargoes from retrograde-transport-mediated depletion at synaptic terminals.

## Results

### EV cargoes are locally depleted at nwk mutant presynaptic terminals

EVs at the *Drosophila* larval neuromuscular junction (NMJ) derive from neuronal membranes, and thus contain neuronal glycoproteins that can be detected using α-HRP antibodies (Fuentes-Medel et al., 2009; Snow et al., 1987; Walsh et al., 2021). During our studies of the endocytic F-BAR and SH3 domain-containing protein Nervous Wreck (Nwk), we found that *nwk* null mutants exhibited a strong reduction in the intensity of postsynaptic α-HRP puncta (within 3 μm of the boundary of motor neurons innervating muscles 6 and 7) (**Fig. 1A**). This suggests that in addition to its previously defined roles in synaptic morphogenesis, growth factor receptor trafficking, endocytosis, and synaptic transmission (Coyle et al., 2004; Del Signore et al., 2021; O’Connor-Giles et al., 2008; Rodal et al., 2008; Ukken et al., 2016), presynaptic Nwk may also regulate EV cargo traffic at synapses.

**Figure 1:**
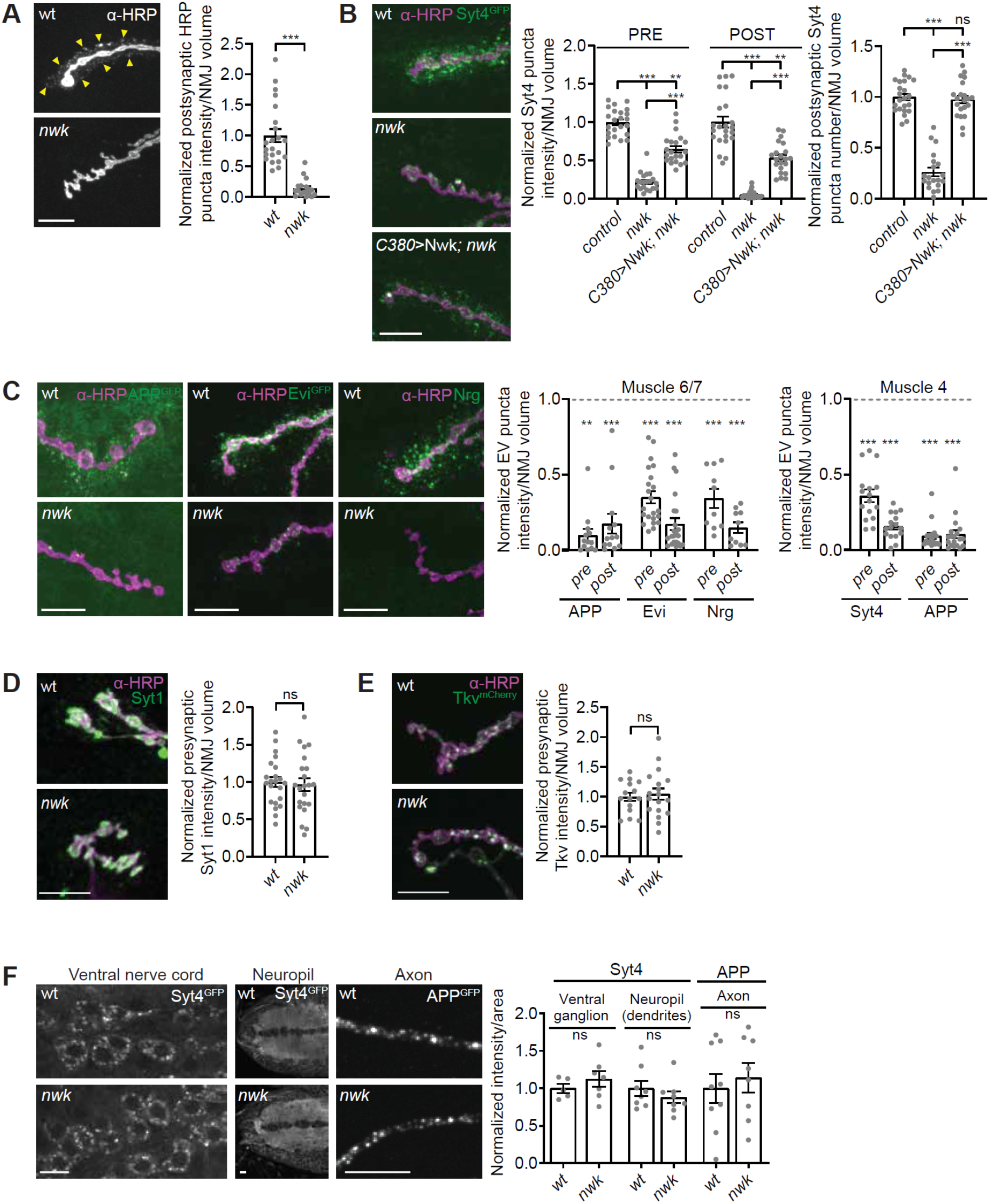
EV cargoes are locally depleted at *nwk* mutant synapses. **(A)** *nwk* mutant synapses lack α-HRP-labeled postsynaptic neuronal membrane puncta (indicated by arrowheads). (Left) Representative images of muscle 6/7 NMJs. (Right) Quantification of postsynaptic α-HRP puncta intensity. **(B)** *nwk* mutants cause neuron-autonomous pre- and post-synaptic depletion of the endogenously tagged EV cargo Syt4-GFP. C380 is a neuron-specific driver. (Left) Representative images of muscle 6/7 NMJs. (Right) Quantification of Syt4-GFP puncta intensity and number. **(C)** EV cargoes APP-GFP, Evi-GFP, Nrg, and Syt4-GFP are reduced at *nwk* mutant synapses. (Left) Representative images of muscle 6/7 NMJs. (Middle) Quantification of EV cargo puncta intensity at muscle 6/7. (Right) Quantification of EV cargo puncta intensity at muscle 4. Dashed line indicates mean EV cargo levels at control NMJs **(D-E)** Levels of the non-EV cargoes Syt1 and Tkv-mCherry are unaffected at *nwk* mutant synapses: (Left) Representative images of muscle 6/7 NMJs (Syt1) and muscle 4 NMJs (Tkv-mCherry). (Right) Quantification of Syt1 or Tkv-mCherry puncta intensity. **(F)** EV cargo levels remain unchanged in cell bodies, neuropil, and axons in *nwk* mutants: (Left) Representative images of Syt4-GFP in ventral ganglion cell bodies and neuropil, and APP-GFP in axons. (Right) Quantification of Syt4-GFP or APP-GFP intensity. Data is represented as mean +/− s.e.m.; n represents NMJs **(A-E)**, or axons or brains **(F)**. NMJ intensity measurements were normalized to presynaptic volume; all measurements were further normalized to the mean of their respective controls. All scale bars are 10 μm. **Associated with Figure S1. See Tables S1 and S3 for detailed genotypes and statistical analyses.**

α-HRP antibodies are likely to detect many neuronal proteins undergoing a variety of trafficking itineraries (Snow et al., 1987), only a small fraction of which result in EV sorting. Therefore, we next tested the role of *nwk* in the traffic of specific established EV cargoes. First, using endogenously tagged Syt4-GFP (Walsh et al., 2021), we found that similar to its α-HRP phenotype, loss of *nwk* led to reduced mean postsynaptic Syt4-GFP levels, together with a decreased number of postsynaptic Syt4-GFP puncta (**Fig 1B**). Importantly, using this single, defined cargo, we also detected a striking reduction presynaptically (**Fig 1B**). These phenotypes were rescued by neuronal re-expression of Nwk using the binary GAL4/UAS system (**Fig 1B**). Taken together, these results suggest that Nwk cell autonomously regulates Syt4 levels in EV donor neurons, resulting in reduced release of Syt4 in EVs. Loss of *nwk* similarly caused a dramatic decrease in pre- and post-synaptic levels of several other known EV cargoes, including neuronal GAL4-driven human Amyloid Precursor Protein (hAPP-EGFP (Walsh et al., 2021)), the neuronal GAL4-driven long isoform of Evenness Interrupted (Evi) tagged with EGFP (EviL-EGFP (Korkut et al., 2009)), and the endogenous neuronal isoform of Neuroglian (Nrg (Walsh et al., 2021)) (**Fig 1C and Supplementary Fig 1A-C**). This phenotype was not specific to muscle 6/7, as we observed a similar effect at the NMJ on muscle 4 (**Fig 1C and Supplementary Fig 1D,E**). Therefore, the effects of *nwk* mutants are consistent for multiple neurons and EV cargoes. These observations highlight a new role for Nwk in regulating the levels of EV cargo proteins in donor synapses, and thus in the recipient postsynaptic cell.

To test if this effect was specific to EV cargoes, we measured the presynaptic levels of two transmembrane proteins that are not normally trafficked into EVs: the calcium sensor Synaptotagmin-1 (Syt1), which is associated with synaptic vesicles (Littleton et al., 1993), and the BMP signaling receptor Thickveins (Tkv), which localizes to the plasma membrane and endosomes (Deshpande et al., 2016; Smith et al., 2012). We found that presynaptic levels of both endogenous Syt1 and neuronally GAL4-driven UAS-Tkv-mCherry were similar between *nwk* mutants and controls (**Fig 1D, E**), suggesting that reduction of Syt4, hAPP, Evi, and Nrg in *nwk* mutants is specific to the EV trafficking itinerary.

We next asked whether Nwk functions to maintain EV cargo levels specifically at synaptic terminals or throughout the neuron. We measured the mean intensity of Syt4-GFP in cell bodies and dendrites within the larval ventral nerve cord, and the mean intensity of hAPP-EGFP (as the Syt4-GFP signal was too faint for this analysis) in the region of the motor neuron axon closest to muscle 4 synaptic terminals. We found that *nwk* mutants had similar levels of Syt4-GFP in cell bodies and dendrites compared to controls, and similar levels of hAPP-EGFP in their axons compared to controls (**Fig 1F**). Together these results suggest that Nwk specifically and locally regulates the levels of EV cargo proteins at synaptic terminals.

### nwk mutants phenocopy loss of EV cargo function

We next asked if depletion of EV cargoes at *nwk* mutant synapses causes a loss of EV cargo function. At the NMJ, traffic of Syt4 into EVs is thought to regulate multiple forms of activity-dependent synaptic growth and plasticity (Barber et al., 2009; Harris et al., 2018; Korkut et al., 2013; Piccioli and Littleton, 2014; Yoshihara et al., 2005). In this system, high frequency stimulation causes an increase in the frequency of miniature excitatory junction potentials (mEJPs). This functional plasticity is termed high-frequency stimulation-induced miniature release (HFMR) and depends on presynaptically-derived Syt4 (Korkut et al., 2013). To directly test whether *nwk* mutants exhibit a loss of Syt4 function and therefore phenocopy *syt4* mutant defects, we measured the responses of *nwk* mutant synapses to high frequency stimulation, and found that HFMR was strongly reduced in *nwk* mutants (**Fig. 2A,B**), similar to *syt4* mutants (Yoshihara et al., 2005). This phenotype could be rescued by neuronal but not muscle GAL4-driven expression of *nwk,* indicating that HFMR specifically requires presynaptic *nwk* (**Fig. 2C**).

**Figure 2:**
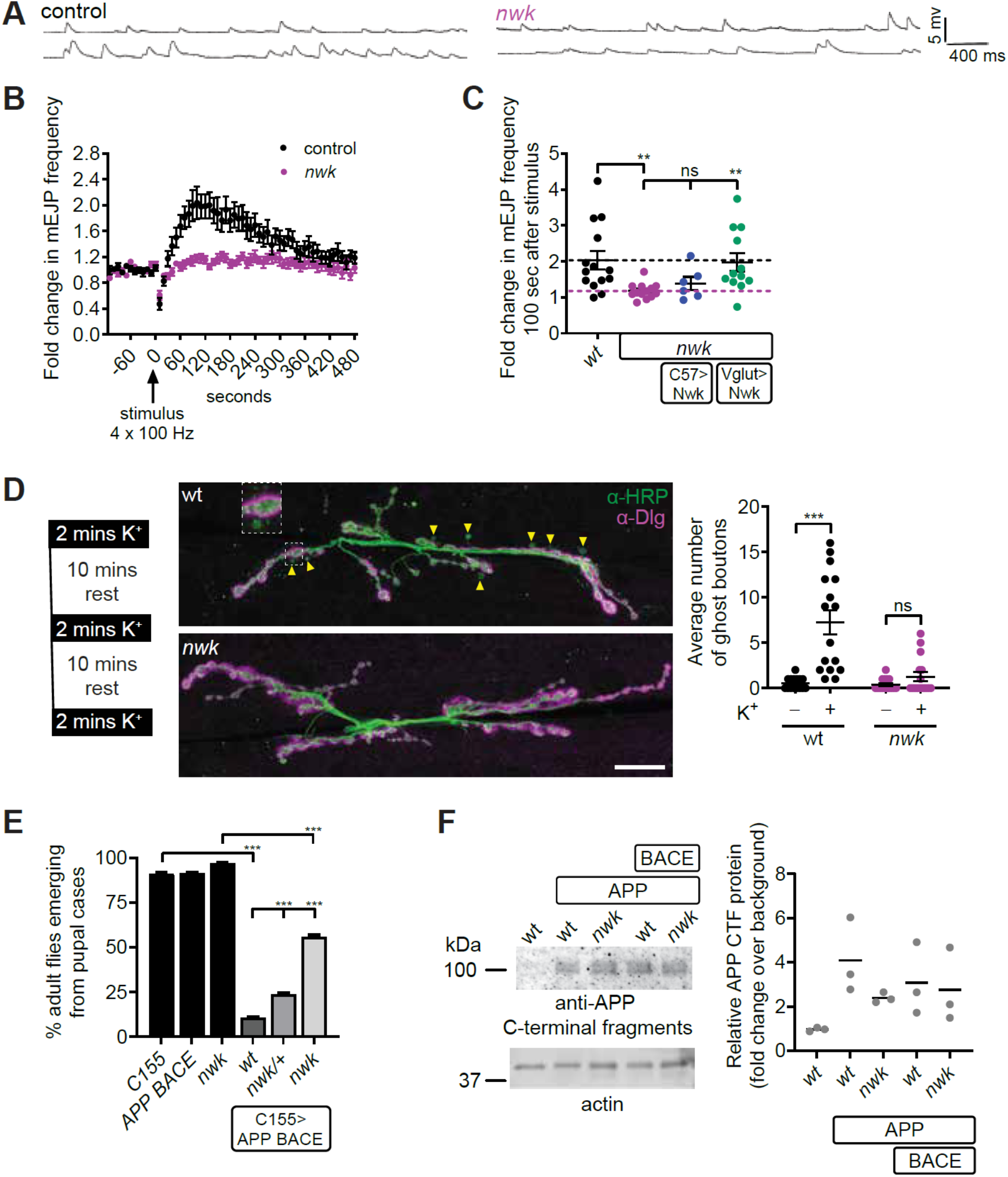
*Nwk* mutants exhibit loss of EV cargo function. **(A-C)** Syt4-dependent functional plasticity (high frequency stimulation-induced mEJP release (HFMR)) is abolished in *nwk* mutants. **(A)** Representative traces of mEJPs before (top trace) and after (bottom trace) high frequency stimulation (4 x 100 Hz). (**B**) Timecourse of mEJP frequency after stimulation. **(C)** *nwk* mutant HFMR phenotype is cell autonomous to neurons. Scatter plot of mEJP frequency 100 seconds after high frequency stimulation. Black dashed line represents *wt* mean value and the pink dashed line represents *nwk* mean value. C57-GAL4 and Vglut-GAL4 are muscle and neuron-specific drivers, respectively. **(D)** Syt4-dependent structural plasticity (ghost bouton budding) is abolished in *nwk* mutants. (Left) Paradigm for spaced stimulation with high K^+^/Ca^2+^. (Middle) Representative maxIPs of confocal stacks at muscle 6/7 labeled with α-HRP (presynaptic) and α-Dlg (postsynaptic antibodies). Arrowheads point to ghost boutons, containing HRP but not Dlg, enlarged in inset. Scale bar is 20 μm. (Right) Quantification of ghost boutons with and without stimulation. **(E)** APP BACE-induced eclosion defect is suppressed in *nwk* mutants. Graph shows the percentage of adult flies emerging from pupal cases. **(F)** GAL4^C155^-driven APP levels are comparable in wt and *nwk* heads: Immunoblot of *Drosophila* head extracts with α-APP C-terminal fragment and α-actin antibodies. Quantification shows fold change of APP relative to actin levels. Data is represented as mean +/−s.e.m.; n represents NMJs (**B-D**), pupal cases (**E**), or biological replicates (**F**). **See Tables S1 and S3 for detailed genotypes and statistical analyses.**

We next tested whether Nwk is required for Syt4 dependent structural plasticity. Spaced potassium stimulation of wild-type *Drosophila* synapses leads to the Syt4-dependent formation of nascent synapses, termed ghost boutons, which contain presynaptic markers but have not yet assembled postsynaptic components such as Discs Large (Dlg) (Ataman et al., 2008; Korkut et al., 2013; Piccioli and Littleton, 2014). In contrast to the significant increase in ghost bouton formation following spaced stimulation in control animals, *nwk* mutants did not show an increase (**Fig 2D**). Together these results indicate that *nwk* mutants exhibit a loss of Syt4 function in activity-dependent synaptic growth and plasticity. Since *nwk* mutants have normal Syt4 levels in cell bodies and neuropil, these results further suggest that Syt4 specifically requires presynaptic localization for its functions.

As loss of *nwk* led to a decrease in hAPP at NMJs (**Fig 1C and Supplementary Fig 1A,E**), we tested if it also reduces hAPP-induced toxicity. GAL4-mediated neuronal overexpression of hAPP and the amyloidogenic protease beta secretase (BACE) leads to defects including decreased eclosion (Chakraborty et al., 2011; Greeve et al., 2004; Mhatre et al., 2014) (**Fig 2E**). Remarkably, this eclosion defect was significantly suppressed in both *nwk* heterozygous and *nwk* homozygous mutant backgrounds, suggesting that loss of even one copy of *nwk* can reduce the toxicity of hAPP and BACE in the adult nervous system (**Fig 2E**). Suppression was not due to broad changes in hAPP levels in adult fly head protein extracts, again suggesting a local and specific effect (**Fig. 2F**). Together these results show that the depletion of EV cargoes Syt4 and hAPP at *nwk* mutant synaptic terminals correlates with a loss of function for these EV cargos in the nervous system.

### Endocytic machinery regulates EV cargo traffic at synapses

Nwk is a conserved membrane-remodeling protein that physically interacts with components of the canonical clathrin-mediated endocytic machinery (Almeida-Souza et al., 2018; Del Signore et al., 2021; O’Connor-Giles et al., 2008; Rodal et al., 2008; Xiao et al., 2018; Xiao and Schmid, 2020). At *Drosophila* synapses, this machinery controls synaptic vesicle recycling, synaptic transmission, and growth factor receptor trafficking (Kaempf and Maritzen, 2017). We hypothesized that like Nwk, other endocytic proteins may also regulate EV cargo traffic at synapses. We therefore investigated EV cargo traffic in mutants of endocytic proteins of diverse functions including membrane lipid composition, membrane remodeling, and cytoskeletal regulation. Loss of the lipid phosphatase synaptojanin (synj) or presynaptic knockdown of the endocytic adapter Dap160/intersectin led to a depletion of Syt4-GFP both pre- and postsynaptically (**Fig 3A and Supplementary Fig 2**). Further, neuronal expression of a dominant negative dynamin mutant (Shi^K44A^) or presynaptic knockdown of the membrane deforming protein Endophilin A caused reduction of postsynapticα-HRP puncta (**Fig 3B and Supplementary Fig 2**). By contrast, both pre- and postsynaptic Syt4-GFP levels were slightly increased in null mutants of the BAR-SH3 sorting nexin SH3PX1/Snx9, though it physically interacts with Nwk (Ukken et al., 2016) (**Fig 3A**). These results indicate that maintenance of EV cargo levels at synapses is a function shared by some but not all Nwk-interacting endocytic machinery.

**Figure 3.**
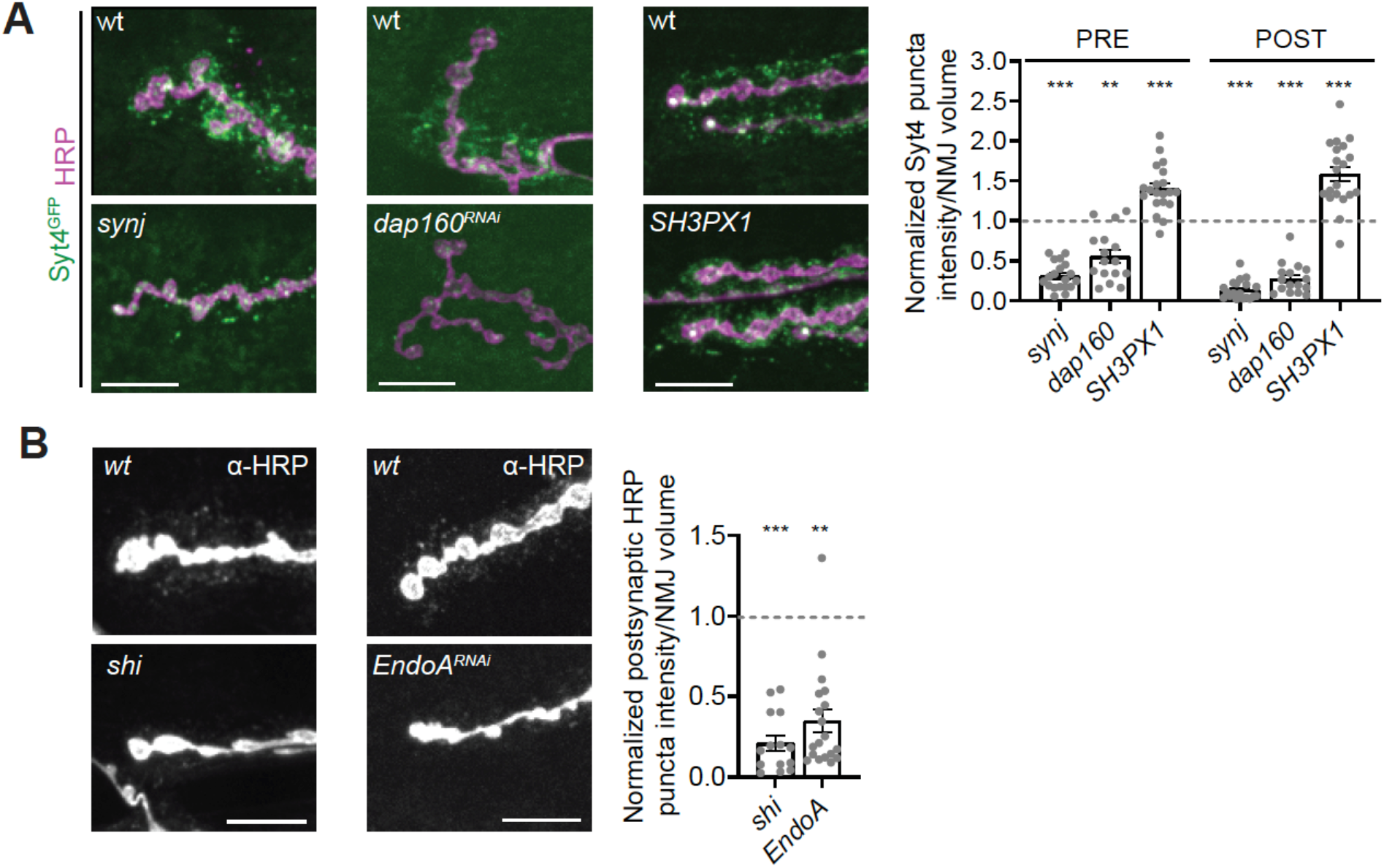
A subset of endocytic machinery is required for EV cargo trafficking. Presynaptic and postsynaptic EV cargo levels are reduced in multiple endocytic mutants. **(A)** (Left) Representative images of muscle 6/7 NMJs. (Right) Quantification of Syt4-GFP puncta intensity. (**B**) (Left) Representative images of α-HRP debris on muscle 6/7 NMJs. (Right) Quantification of α-HRP debris intensity. Data is represented as mean +/− s.e.m.; n represents NMJs. NMJ intensity measurements were normalized to presynaptic volume; all measurements were further normalized to the mean of their respective controls, which is indicated with a dashed line. All scale bars are 10 μm. **Associated with Figure S2. See Tables S1 and S3 for detailed genotypes and statistical analyses.**

The canonical function of endocytic proteins is to control clathrin-mediated endocytosis, which is the primary mechanism for synaptic vesicle recycling at the *Drosophila* larval NMJ (Heerssen et al., 2008; Kasprowicz et al., 2008), and is required for synaptic morphogenesis via endocytic traffic of growth factor receptors (Choudhury et al., 2016; Dwivedi et al., 2021). We asked if clathrin and its heterotetrameric AP-2 adaptor complex were similarly involved in synaptic EV cargo traffic. We found that mutants lacking the AP-2σ subunit exhibited a significant decrease in pre- and postsynaptic levels of Syt4-GFP (**Fig 4A, Supplementary Fig 3A-B**). Surprisingly, we found mutants lacking either the AP-2μ or the AP-2α subunit exhibited only a mild decrease in presynaptic levels of Syt4-GFP, and no significant change postsynaptically (**Fig 4A, Supplementary Fig 3A-B**). This suggests that the levels of EV cargo at synapses specifically depend on the AP-2σ subunit, or that there is redundancy between the AP-2α or AP-2μ subunits, as has been previously suggested (Gu et al., 2013). To decipher between these possibilities, we measured Syt4-GFP levels in *AP-2α; AP-2μ* double mutants and found a striking decrease in pre- and postsynaptic levels of Syt4-GFP, similar to *nwk* mutants (**Fig. 4A, Supplementary Fig 3A-B**). Thus, EV cargo levels are dependent on the AP-2 clathrin adaptor complex with functional redundancies among the AP-2α and AP-2μ subunits.

**Figure 4.**
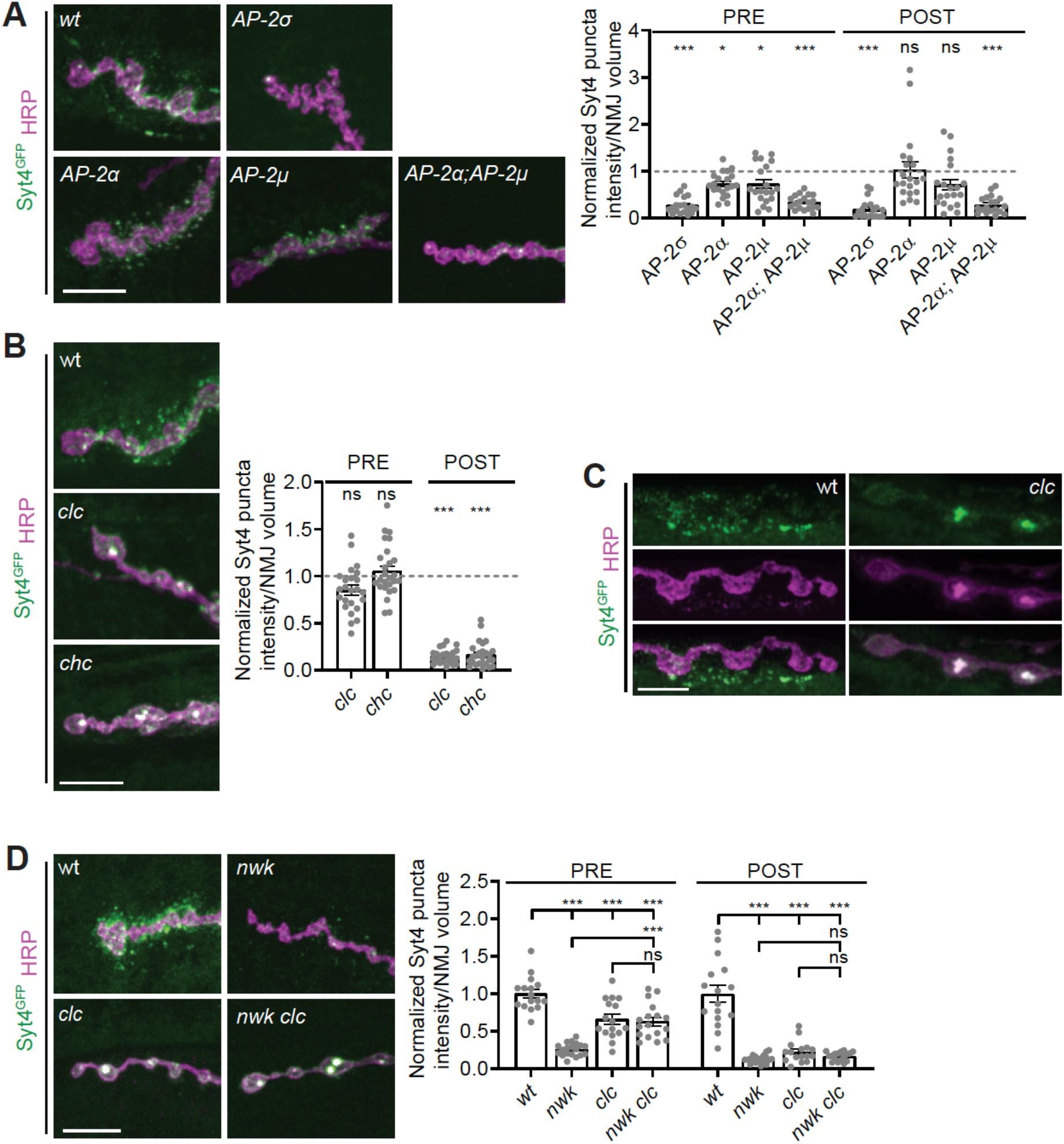
EV cargo trafficking is clathrin-dependent. **(A)** Presynaptic and postsynaptic EV cargo levels are reduced in clathrin adaptor AP2 mutants. (Left) Representative images of muscle 6/7 NMJs. (Right) Quantification of Syt4-GFP puncta intensity. **(B)** EV cargoes show altered localization in clathrin mutants and are not released from neurons. (Left) Representative images of muscle 6/7 NMJs. (Right) Quantification of Syt4-GFP puncta intensity. **(C)** Airyscan microscopy of *clc* mutant muscle 6/7 NMJs show that EV cargo accumulations extend into the interior of the bouton. **(D)** EV cargo accumulation in clathrin mutants is epistatic to its depletion in *nwk* mutants. (Left) Representative images of muscle 6/7 NMJs. (Right) Quantification of Syt4-GFP puncta intensity. Data is represented as mean +/− s.e.m.; n represents NMJs. NMJ intensity measurements were normalized to presynaptic volume; all measurements were further normalized to the mean of their respective controls, which is indicated with a dashed line. All scale bars are 10 μm. **Associated with Figure S3. See Tables S1 and S3 for detailed genotypes and statistical analyses.**

The importance of the AP-2 clathrin adaptor complex in EV cargo regulation prompted us to test directly whether EV cargo levels at synapses are dependent on clathrin itself. We examined Syt4-GFP in hypomorphic mutants of clathrin light chain (*clc*) or clathrin heavy chain (*chc*). We found no change in presynaptic levels of Syt4-GFP in *clc* and *chc* mutants compared to controls, but its localization was drastically shifted into largeα-HRP-positive presynaptic accumulations (**Fig. 4B,C, Supplementary Fig 3C-D**). This change in Sy4-GFP presynaptic localization in *clc* and *chc* mutants was associated with a striking decrease in postsynaptic levels of Syt4-GFP compared to controls (**Fig. 4B, Supplementary Fig 3C-D**), suggesting that Syt4-GFP is not released in EVs from clathrin mutant synapses. Inactivation of clathrin is known to lead to the formation of large cisternal membrane compartments derived from bulk membrane uptake in synapses (Heerssen et al., 2008; Kasprowicz et al., 2008), which may correspond to the Syt4-GFP and HRP-positive accumulations that we see in *clc* and *chc* synapses. As *nwk* mutants exhibit a significant depletion of EV cargoes both pre- and postsynaptically (**Fig. 1B,C**), we wondered whether these presynaptic accumulations of Syt4-GFP in *clc* and *chc* mutants would be sensitive or resistant to loss of *nwk*. We found that simultaneous loss of *clc* and *nwk* phenocopied loss of *clc* alone with regards to levels and localization of Syt4-GFP pre- and postsynaptically (**Fig. 4D**). This suggests that the Syt4-GFP accumulations in *clc* mutant synapses are not accessible to the mechanism by which EV cargoes are depleted at *nwk* mutant synapses. Together these results highlight a novel clathrin-dependent mechanism that regulates the traffic and levels of EV cargoes at synapses.

### Nwk opposes the function of Vps35/retromer in EV cargo trafficking

We next explored the mechanisms by which Nwk-associated membrane-remodeling machinery may control the distribution of EV cargoes between release in EVs versus synaptic depletion. One possibility is that this machinery acts at the plasma membrane in a non-canonical role, to regulate fusion of EV-containing MVBs (similar to proposed functions of dynamin at chromaffin granule fusion pores (Shin et al., 2018)) and that in its absence these MVBs are targeted for degradation. Alternatively, endocytic machinery could serve a canonical function in endocytic loading of cargoes into the EV-permissive precursor compartment that populates MVBs destined for EV release. Mutation of the retromer complex component *Vps35* causes accumulation of EV cargoes in these precursor compartments and a concomitant increased release of EVs (Walsh et al., 2021). If the Nwk-dependent endocytic machinery acts to promote EV fusion with the plasma membrane, we would expect to see recovery of presynaptic cargo levels in *Vps35*, *nwk* double mutants relative to *nwk* single mutants, but no postsynaptic recovery, as MVBs would not be released (**Fig 5A, “release”**). On the other hand, if Nwk plays a more canonical role in loading MVB precursors prior to their release, we would expect to see recovery of both presynaptic and postsynaptic EV cargo levels in *Vps35*, *nwk* mutants relative to *nwk* single mutants (**Fig 5A, “sorting”**). We found that loss of *Vps35* in the *nwk* mutant background significantly rescued Syt4-GFP levels both pre- and postsynaptically, and also significantly restored the number of postsynaptic Syt4-GFP puncta (**Fig. 5B**), suggesting that Nwk is involved in loading EV precursors and maintaining presynaptic cargo levels, rather than in fusion of MVBs and release of EVs from the neuron.

**Figure 5:**
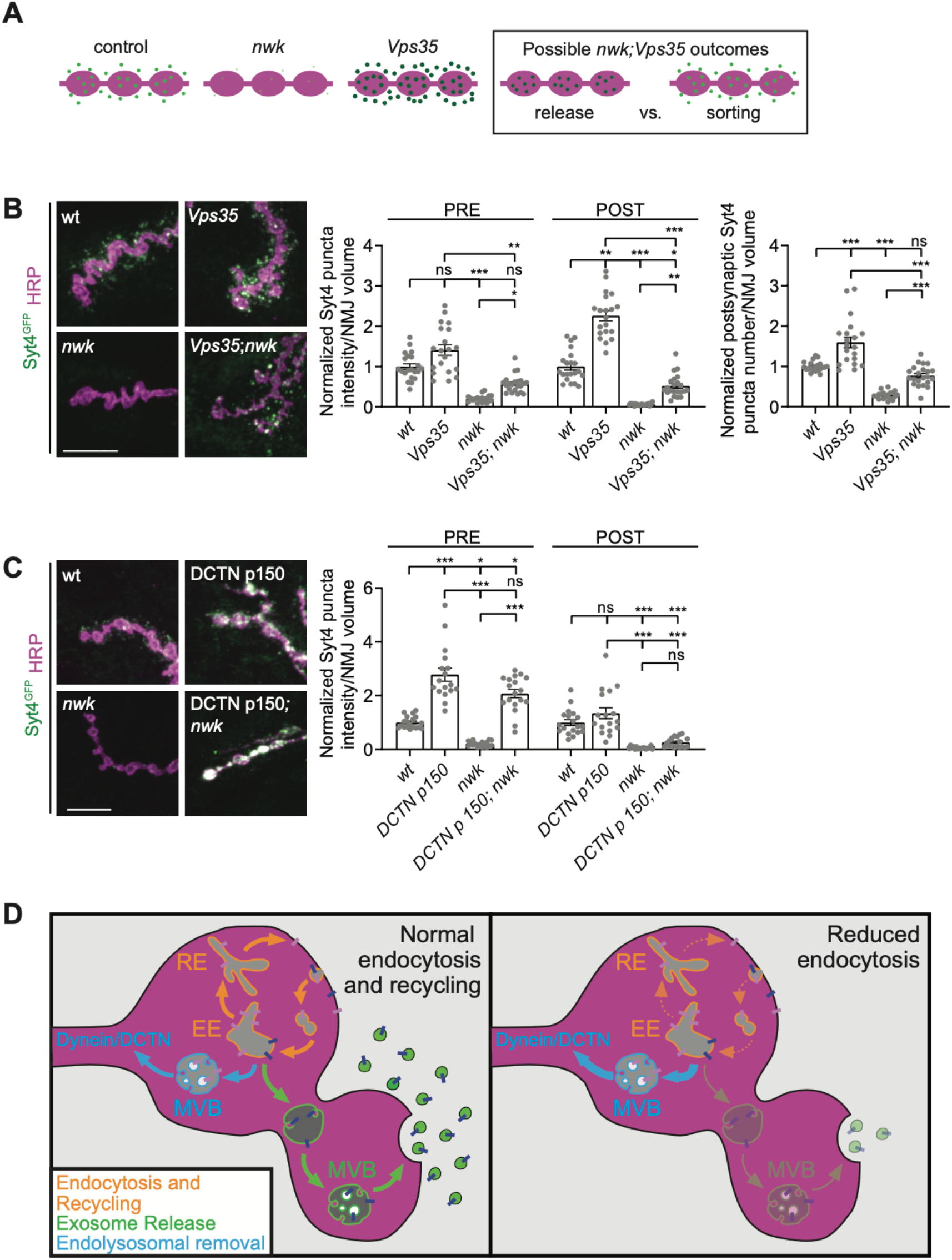
Nwk opposes retromer in loading EV precursors, and acts upstream of dynactin-mediated retrograde transport. **(A)** Cartoon depicting potential EV phenotype outcomes for *nwk; Vps35* experiment, testing if Nwk regulates MVB-PM release or EV cargo sorting. **(B)** EV cargoes return to normal levels both pre- and post-synaptically in *nwk; Vps35* double mutants (Left) Representative images of muscle 6/7 NMJs. (Right) Quantification of Syt4-GFP puncta intensity and number. **(C)** EV cargoes are retained at the synapse upon p150-DCTN overexpression, but these compartments are not competent for EV release in the *nwk* mutant background. (Left) Representative images of muscle 6/7 NMJs. (Right) Quantification of Syt4-GFP puncta intensity. **(D)** At wild type synapses, proper traffic and function of EV cargoes depends on a robust flux through the recycling pathway (left, orange arrows). However, endocytic mutant synapses exhibit a depletion of EV cargoes, likely due to reduced recycling flux, resulting in aberrant sorting for degradation upstream of dynactin/dynein mediated retrograde transport (right). Data is represented as mean +/− s.e.m.; n represents NMJs. NMJ intensity measurements were normalized to presynaptic volume; all measurements were further normalized to the mean of wild type controls. All scale bars are 10 μm. **See Tables S1 and S3 for detailed genotypes and statistical analyses.**

### Nwk acts upstream of dynactin-mediated retrograde transport

Our data suggest that endocytic machinery protects EV cargoes from degradation by loading them into an EV-permissive precursor compartment. We hypothesized that in *nwk* mutants, these EV cargoes may be depleted from the synapse via retrograde traffic to the cell body for lysosomal degradation. To test this, we overexpressed a dominant negative dynactin (DCTN) truncated p150/Glued subunit to inhibit dynein/dynactin-mediated retrograde transport (Allen et al., 1999). In otherwise wild-type synapses expressing DCTN-p150^Δ^, we found that presynaptic Syt4-GFP levels more than doubled (**Fig. 5C**), indicating that EV cargoes do undergo retrograde transport, and build up at the synapse in the absence of dynein function. Interestingly, there was no concomitant increase in postsynaptic Syt4-EGFP levels (**Fig. 5C**), indicating that the cargoes destined for retrograde transport are located in a non-EV permissive compartment. We then used this system to test whether in *nwk* mutants EV cargoes are depleted from the synapse due to retrograde transport. We overexpressed DCTN-p150^Δ^ in the *nwk* mutant background and found the same striking increase in presynaptic Syt4-GFP levels as in wild-type synapses (**Fig. 5C**), suggesting that in *nwk* mutants EV cargoes are largely being targeted for retrograde transport. Further, postsynaptic Syt4-GFP levels remained depleted relative to wild type controls (**Fig. 5C**), indicating that in *nwk* mutants synaptic EV cargoes that are destined for retrograde transport are in a non-EV permissive compartment. Overall, our data suggest that synaptic depletion of EV cargoes in *nwk* mutants results from a shift of cargoes towards a compartment that cannot produce EVs, and is sent to the cell body via retrograde transport (**Fig 5D**).

## Discussion

Here we report a novel function of clathrin-mediated endocytic machinery in regulating trafficking of EV cargoes at synapses *in vivo*. We find that endocytic mutants exhibit a local depletion of EV cargoes at synaptic terminals, correlating with a loss of Syt4-dependent synaptic plasticity at the larval NMJ and a reduction in APP-dependent toxicity in the adult nervous system. Mechanistically, endocytic machinery opposes the retromer complex in sorting cargoes into EV-precursor compartments and acts upstream of dynactin-mediated retrograde transport. Together our data support a model in which endocytic machinery protects EV cargoes from local depletion at synaptic terminals, consequently preserving cargo function and promoting its release in EVs. Our results suggest a new interpretation for previously reported phenotypes of endocytic mutants and uncover a new pathway for investigating and therapeutically intervening in EV traffic.

### EV traffic is genetically separable from canonical functions of the endocytic machinery

Our new findings add to the diverse functions of the endocytic machinery in neurons, including synaptic vesicle recycling, release site clearance, synaptic transmission, signaling receptor trafficking, and synaptic growth (Chanaday et al., 2019; Deshpande and Rodal, 2015). If EV cargo trafficking defects in endocytic mutants were indirectly caused by impairment of these canonical roles, then we would expect the severity of EV phenotypes to consistently scale with defects in these other functions. However, we find that mutants with more severe defects in synaptic vesicle recycling (e.g. *AP-2α*, (Gonzalez-Gaitan and Jackle, 1997)) exhibit very mild EV phenotypes, while mutants with mild synaptic vesicle cycling phenotypes (e.g. *nwk* and *dap160* (Del Signore et al., 2021; Koh et al., 2004; Marie et al., 2004)) have severe EV defects. In a similar vein, *AP-2α*, *AP-2μ* and *dap160* mutants exhibit severe synaptic growth phenotypes (Choudhury et al., 2016; Dwivedi et al., 2021; Koh et al., 2004; Marie et al., 2004), likely due to failure to downregulate BMP (Bone Morphogenetic Protein) receptor signaling (Deshpande and Rodal, 2015). Among these, only *dap160* has a severe EV defect. By contrast, *nwk* mutants have a relatively mild synaptic growth phenotype (Coyle et al., 2004) and a severe EV phenotype. *Vps35/*retromer mutants also exhibit BMP-pathway dependent synaptic overgrowth (Korolchuk et al., 2007), but conversely accumulate excess EV cargoes rather than depleting them (Walsh et al., 2021). Thus, synaptic growth is not functionally coupled to EV cargo traffic, levels or release. Finally, *nwk* and *SH3PX1* share a role in synaptic transmission (Coyle et al., 2004; Ukken et al., 2016). However, this role is not linked to their EV phenotypes, as we found that *SH3PX1* mutants exhibit a significant increase in EV cargo levels, opposite to *nwk* mutants. Overall, our results indicate that the role of endocytic machinery in regulating EV cargo traffic is genetically separable from its other functions in the nervous system, and instead reflects an independent and novel function of these conserved molecules at synaptic terminals.

### Endocytic machinery sorts cargoes into EV-precursor compartments

What then is the mechanism by which endocytic machinery regulates EV cargo levels? Our data indicate that Nwk functions in an opposing pathway to the retromer complex, to load cargoes into EV-permissive compartments. One possibility is that this occurs via the canonical role of the clathrin/AP-2 endocytic machinery at the plasma membrane (Kaksonen and Roux, 2018). Our data differentiating the phenotypes of clathrin and AP-2 mutants provides some insights into the specific mechanisms involved: We find that clathrin is strictly required to form functional EV precursors, and in its absence, cargoes accumulate in large cisternae. By contrast, loss of AP-2 adaptors and other endocytic machinery leads to cargo depletion, perhaps due to a slowed but still partially functional endocytic pathway (Chen and Schmid, 2020; Dickman et al., 2005). Indeed, efficient clathrin-mediated endocytic internalization could be required for effective EV loading, by increasing the flux of cargo through the endosome-plasma membrane recycling pathway that populates these EV precursor compartments ((Walsh et al., 2021), **Fig. 5D**, orange arrows). In support of this hypothesis, loss of *nwk* phenocopies mutation of the recycling endosome regulator Rab11, including presynaptic depletion of EV cargoes in opposition to retromer, and loss of Syt4 physiological function (Ashley et al., 2018; Koles et al., 2012; Korkut et al., 2013; Walsh et al., 2021). Finally, our observation that AP2- and AP2-μ exhibit partial redundancy indicates that hemicomplexes of AP-2 may be sufficient to sort EV cargo (Gu et al., 2013), or that other adaptors such as AP180 may also function in this pathway (Bao et al., 2005).

Alternatively, clathrin-mediated endocytic machinery may be acting in a noncanonical role on intracellular compartments to sort EV cargoes. Clathrin-mediated endocytic components are required to reform synaptic vesicles from endosomal intermediates, particularly during elevated activity (Heerssen et al., 2008; Kasprowicz et al., 2008; Kasprowicz et al., 2014; Kittelmann et al., 2013; Kononenko et al., 2014; Watanabe et al., 2014). Clathrin localizes directly to endosomes, where it regulates endosomal cargo sorting and intralumenal vesicle formation (Raiborg et al., 2001; Raiborg et al., 2006; Sachse et al., 2002; Shi et al., 2009; Wenzel et al., 2018). Further, the mammalian Nwk homolog FCHSD2, in addition to its role at the internalization step of endocytosis (Almeida-Souza et al., 2018; Xiao et al., 2018), suppresses recycling and lysosomal degradation of signaling receptors (Xiao and Schmid, 2020). The extreme density of endocytic machinery at synaptic membranes makes it difficult to detect a small endosomally localized fraction of this machinery, though Nwk does exhibit partial co-localization with Rab11 (Rodal et al., 2008). Future studies will be necessary to discriminate between whether clathrin-mediated endocytic machinery indirectly loads EV-precursor endosomes via canonical endocytosis, or if it instead acts on them directly.

### EV cargo sorting for release occurs upstream of retrograde transport

Our data also provides new insight into the overall trafficking itineraries of neuronal EV cargoes. Using a dynactin mutant, we identified a population of EV cargo-containing compartments that retrogradely traffics from synaptic terminals. Trapping these compartments at *nwk* mutant synapses is sufficient to restore presynaptic levels of EV cargoes, but does not allow release of these cargoes from the neuron. Thus, Nwk-mediated cargo sorting occurs upstream of retrograde transport, and the missorted cargoes in the *nwk* mutant are no longer EV-competent and are destined for synaptic removal, likely for degradation in the cell body. Degradative lysosomes are enriched in cell bodies relative to axons or at synaptic terminals (Ferguson, 2018; Lie et al., 2021), and there is also evidence of local protein degradation along axons or at axon terminals (Farfel-Becker et al., 2019; Jin et al., 2018). Our data suggest for the first time that synaptic EV cargoes are cleared from the synapse after transport rather than on-site degradation, though we cannot exclude the possibility of separate pools of cargoes: some that are locally degraded, and some that are targeted for retrograde transport. Future *in vivo* investigations focused on the role of retrograde trafficking machinery and their adaptors may yield insight into additional points by which neurons control release versus degradation of EV cargoes at distinct subcellular locations (Heisler et al., 2018; van Niel et al., 2018). Interestingly, the phenotypes we observed for endocytic mutants (reduction in cargo levels) and retromer (increase in cargo levels) at mature synapses are opposite to what has been found for EV-relevant cargoes in other cell types. For example, in *C. elegans* Wntless/Evi accumulates near the plasma membrane in *dpy-23*/AP-2 mutants and is depleted in *vps-35* mutants (Pan et al., 2008; Yang et al., 2008). This highlights the cell type specificity of trafficking trajectories, and the unique cell biology of neurons, likely due to their extreme morphology and limited lysosomal capacity (Ferguson, 2018).

### Implications for the functions of neuronal EVs and for EV-directed disease therapies

While there has been an explosion of research identifying the contents of EVs and the diverse functions of cargoes, the field has been lacking approaches to specifically manipulate their biogenesis and release. Our findings provide new tools to ask where EV cargoes function in the neuron, and to locally manipulate EV cargo levels at synapses. For instance, we found that loss of *nwk* leads to a local decrease in EV cargo levels both pre- and post-synaptically at synaptic terminals, with a corresponding loss of EV cargo function, suggesting that levels of EV cargoes at synaptic terminals (but not elsewhere in the neuron) correlate with EV cargo function. We report that clathrin mutants have normal levels but aberrant localization of presynaptic EV cargoes, leading to a severe reduction in postsynaptic EV cargo levels. These mutants and genetic manipulations provide ways to specifically alter presynaptic and/or postsynaptic levels of EV cargoes, which will be useful for future *in vivo* EV cargo trafficking and functional studies.

Given that local depletion of EV cargoes at *nwk* mutant synapses correlates with a loss of their physiological functions, EV cargo misregulation may also play unrecognized roles in the previously reported phenotypes of endocytic mutants. These may therefore need to be reinterpreted in the context of local synaptic loss of a multitude of EV cargoes (Holm et al., 2018). For example, *Shi*^ts^ mutants have been used in numerous *Drosophila* circuit-mapping studies to block synaptic transmission at the restrictive temperature (Kitamoto, 2001); however these mutants also show defects at the permissive temperature (Dickman et al., 2006), which could inadvertently cause chronic loss of EV cargoes important for that circuit. Further, neurological phenotypes of mice mutant for endocytic machinery could be consistent with EV signaling defects (Holm et al., 2018; Malakooti et al., 2020; Milosevic et al., 2011).

Finally, our results suggest new interpretations of neurological disease mechanism. Defects in EV and endosomal trafficking are linked to neurodegenerative diseases such as Alzheimer’s Disease (Becot et al., 2020; Song et al., 2020; Winckler et al., 2018). Aged neurons exhibit increased APP endocytosis, leading to endosomal dysfunction and synapse loss (Burrinha et al., 2021). Our data predict that this may also promote APP loading into the EV pathway. This is consistent with previous reports suggesting that upregulation of endocytic machinery exacerbates Alzheimer’s Disease phenotypes (Keating et al., 2006; Ren et al., 2008; Yu et al., 2018), and that loss of endocytic machinery can suppress these phenotypes (Zhu et al., 2013). Downregulation of APP, either genetically or by enhancing lysosomal biogenesis, can suppress generation of Aβ, and amyloid plaques in Alzheimer’s Disease (Hung et al., 2021; Xiao et al., 2015). We report a new and synapse-specific mechanism for targeting APP: loss of *nwk* severely reduces synaptic levels of APP and ameliorates its toxicity in the nervous system, while causing only mild defects in the synaptic vesicle cycle and synaptic growth. Our data suggest that therapeutically targeting EV-sensitive components of endocytic machinery could be a strategy to reduce pathological synaptic EV cargoes in neurological disease.

## Materials and Methods

### Drosophila *culture*

Flies were cultured using standard media and techniques. Flies used for experiments were maintained at 25°C, except for RNAi experiments (Dap160 and EndoA) which were maintained at 29°C, and the *chc*^*B*^ and *clc nwk* experiments which were maintained at 20°C. For detailed information on fly stocks used, see **Table S1**, and for detailed genotype information for each figure panel, see **Table S3**.

### Immunohistochemistry

Wandering third instar larvae from density-controlled crosses were dissected in HL3.1 (Feng et al., 2004) and fixed in HL3.1 with 4% paraformaldehyde for 10 minutes. For detailed information on antibodies used in this study, see **Table S2**. Washes and antibody dilutions were conducted using PBS containing 0.2% Triton X-100 (0.2% PBX). Primary antibody incubations were conducted overnight at 4°C, and secondary antibody incubations for 1-2 hours at room temperature. α-HRP incubations were conducted either overnight at 4°C or for 1-2 hours at room temperature. Prior to imaging, fillets were mounted on slides with Vectashield (Vector Labs). α-GFP nanobodies (Nanotag Biotechnologies) were used to amplify Syt4-GFP signal only for **Fig 4C**.

### Image acquisition and analysis

#### Image acquisition

For analysis of EV and non-EV cargoes at the NMJ, muscle 6/7 (segments A2 and A3) and m4 (segments A2, A3, and A4) were imaged at room temperature using Nikon Elements AR software. Z-stacks were acquired using a Nikon Ni-E upright microscope equipped with a 60X (n.a. 1.4) oil immersion objective, a Yokogawa CSU-W1 spinning disk head, and an Andor iXon 897U EMCCD camera. The images in **Fig 4C** were acquired at room temperature with Zen Blue software on a Zeiss LSM880 Fast Airyscan microscope in super resolution acquisition mode, using a 63X (n.a. 1.4) oil immersion objective. Image acquisition settings were identical for all images in each independent experiment.

#### Quantification of presynaptic and postsynaptic EV cargo levels at the NMJ

3D volumetric analyses of presynaptic and postsynaptic EV cargoes at NMJs were carried out using Volocity 6.0 software (Perkin Elmer). For each NMJ image, both type 1s and 1b bouton were retained for analysis while axons were cropped out. Presynaptic volume was determined through manual thresholding to the α-HRP signal (excluding objects smaller than 7 μm^3^ and closing holes), with a 3.3 μm dilation around this signal delineating the postsynaptic volume. Manual thresholding of the EV cargo signal was conducted to ensure measurements included only EV cargo signal above background muscle fluorescence. Postsynaptic objects smaller than 0.015 μm were excluded. EV cargo sum intensity measurements were calculated within these pre- and postsynaptic volumes and normalized to the presynaptic volume. Puncta number was quantified only within the postsynaptic volume.

#### Quantification of Syt4-GFP in cell bodies

This analysis was conducted in FIJI. Syt4-GFP intensity in even-skipped positive motor neurons was measured from a single middle slice through each cell body. Mean Syt4-GFP intensity was calculated by subtracting the mean intensity of an area of background outside of the cell body in the same slice from the mean intensity within the manually selected cell body region. Each data point corresponds to the average mean Syt4-GFP intensity across 8 cell bodies per brain.

#### Quantification of Syt4-GFP levels in neuropil

This analysis was conducted in FIJI. Using single slices through the center of the ventral ganglion, a region within the neuropil was manually selected. Mean Syt4-GFP intensity within this region of neuropil was calculated by subtracting the mean intensity of an area of background outside of the ventral ganglion from the mean intensity within the selected neuropil region.

#### Quantification of APP-EGFP levels in axons

APP-EGFP levels were measured using FIJI in the m4 axon region between the segmental axon bundle and synaptic boutons. Average intensity projections were manually cropped to exclude everything aside from the axon segment to be measured. Manual thresholding to the HRP signal was conducted to determine the region for measurement. To calculate mean APP-EGFP intensity in the axon, the mean intensity of three areas of background per image were averaged and subtracted from the mean APP-EGFP intensity within the HRP-thresholded region.

### Western blot

Heads (15 pooled per genotype) from *Drosophila* aged 3-10 days were homogenized in 50 μl 2x Laemmli buffer. 15ul of extract was loaded in each lane and fractionated by SDS/PAGE and immunoblotted with α-APP-CTF (A8717, Sigma) and α-actin (JLA-20, DHSB). Blots were visualized using a Biorad Chemidoc system and quantified using FIJI with APP-CTF values normalized to the corresponding actin loading control.

### Eclosion assay

Density-controlled crosses were grown at 25°C for 17 days and the number of eclosed and uneclosed pupal cases were counted for each genotype. Eclosion rate was calculated as the number of pupal cases eclosed divided by the total number of pupal cases.

### Ghost bouton budding

Ghost bouton budding experiments were carried out as previously described (Piccioli and Littleton, 2014), with the modification that only muscle 6/7 NMJs from segment 3 were included in the analysis.

### Electrophysiology

Wandering third instar larvae were dissected in HL3 saline (Stewart et al., 1994). Recordings were taken using an AxoClamp 2B amplifier (Axon Instruments, Burlingame, CA). A recording electrode was filled with 3M KCl and inserted into muscle 6 at abdominal segments A3 or A4. A stimulating electrode filled with saline was used to stimulate the severed segmental nerve using an isolated pulse stimulator (2100; A-M Systems). HFMR was induced by four trains of 100 Hz stimuli spaced 2 s apart in 0.3 mM extracellular Ca2+. Miniature excitatory junction potentials (minis) were recorded 2 min before and 10 min after HFMR induction. Mini frequency at indicated time points was calculated in 10-s bins. Fold enhancement was calculated by normalizing to the baseline mini frequency recorded prior to HFMR induction. Analyses were performed using Clampfit 10.0 software (Molecular Devices, Sunnyvale, CA) and Mini Analysis 6.0.3 (Synaptosoft, Inc.). Each n value represents a single muscle recording, with data generated from at least six individual larvae of each genotype arising from at least two independent crosses. Resting membrane potentials were between −50 mV and −75 mV and were not different between genotypes. Input resistances were between 5 MΩ and 10 MΩ and were not different between genotypes.

### Statistical analyses

All graphing and statistical analyses were completed using GraphPad Prism. Datasets were first analyzed with the D’Agostino-Pearson normality test. Normally distributed datasets were analyzed with either an unpaired t-test (two groups) or a one-way ANOVA with Tukey’s multiple comparisons (more than two groups), while not normally distributed datasets were analyzed with either a Mann-Whitney test (two groups) or a Kruskal-Wallis test with Dunn’s multiple comparisons (more than two groups). Error bars report +/− standard error of the mean. The eclosion experiment in **Fig. 2E** was analyzed using a chi-squared test; error bars report standard error of the proportion. Detailed information about statistical analyses performed for each dataset can be found in **Table S3**. * p<0.05, ** p< 0.01, *** p<0.001.

## Acknowledgments

We thank the Bloomington *Drosophila* Stock Center (NIH P40OD018537), the Developmental Studies Hybridoma Bank created by the NICHD of the NIH. We thank the Bloomington *Drosophila* Stock Center (Indiana University, Bloomington, IN, NIH P40OD018537), and the Developmental Studies Hybridoma Bank created by the NICHD of the NIH. We thank Kate O’Connor-Giles and Vivian Budnik for fly lines, and Sultana Bhuiyan, Steven Del Signore, Erica Dresselhaus, Jack Cheng, Biljana Ermanoska, and Matthew Pescosolido for technical assistance and helpful discussions. This work was supported by NINDS grants R01 NS103967 to A.A.R. and F32 NS110123 to C.R.B, T32 MH019929 to A.S., and by Natural Sciences and Engineering Research Council of Canada (NSERC) to B.A.S. (RGPIN-06004). The authors declare no competing financial interests.

