## Supplemental figures and tables for "Local regulation of extracellular vesicle traffic by the synaptic endocytic machinery"

Blanchette et al.

**3 Supplementary Figures with legends**

**Table S1: Fly strains**

**Table S2: Antibodies**

**Table S3: Statistics by Dataset**

### Supplemental Figure #1

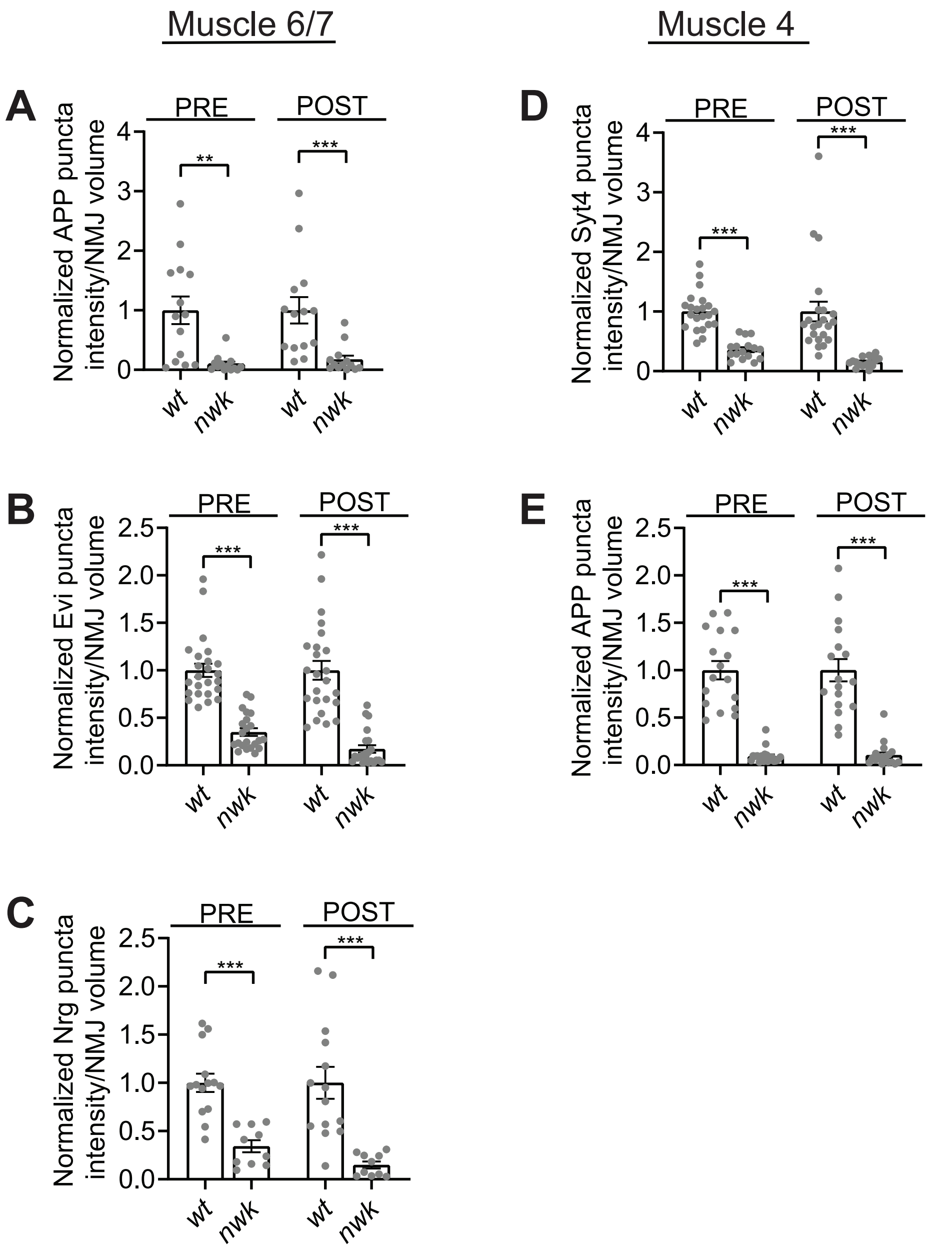


**Figure S1: EV cargo levels are reduced at *nwk* mutant synaptic terminals.**
Fig 1C datasets shown with corresponding controls. (A) Quantification of APP-GFP pre- and postsynaptic puncta intensity at muscle 6/7 NMJs. (B) Quantification of Evi-GFP pre- and postsynaptic puncta intensity at muscle 6/7 NMJs. (C) Quantification of Nrg pre- and postsynaptic puncta intensity muscle 6/7 NMJs. (D) Quantification of Syt4-GFP pre- and postsynaptic puncta intensity at muscle 4 NMJs. (E) Quantification of APP-GFP pre- and postsynaptic puncta intensity at muscle 4 NMJs. Data is represented as mean +/- s.e.m.; n represents NMJs. NMJ intensity measurements were normalized to presynaptic volume; all measurements were further normalized to the mean of their respective controls. **Associated with Figure 1. See Tables S1 and S3 for detailed genotypes and statistical analyses.**

### Supplemental Figure #2

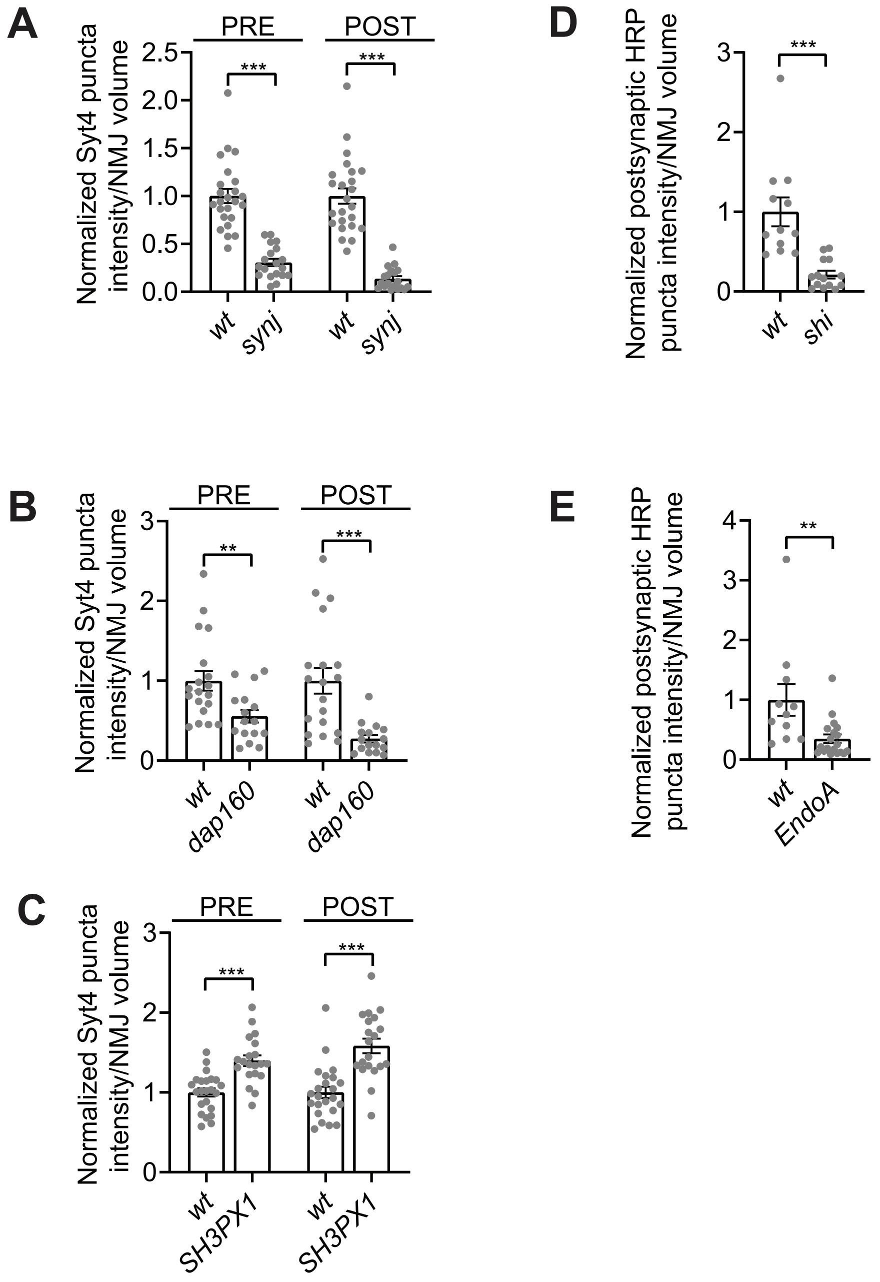


**Figure S2: A subset of endocytic machinery is required for synaptic EV traffic.**

Fig 3A,B datasets shown with corresponding controls. (A) Quantification of pre- and postsynaptic Syt4-GFP puncta intensity in control and *synj* mutants. (B) Quantification of pre- and postsynaptic Sy4-GFP puncta intensity in control and upon neuronal knockdown of *dap160*. (C) Quantification of pre- and postsynaptic Syt4-GFP puncta intensity in control and *SH3PX1* mutants. (D) Quantification of postsynaptic HRP puncta intensity in control and upon neuronal expression of $Shi^{K44A}$. (E) Quantification of postsynaptic HRP puncta intensity in control and upon neuronal knockdown of *EndoA.*

Data is represented as mean +/- s.e.m.; n represents NMJs. NMJ intensity measurements were normalized to presynaptic volume; all measurements were further normalized to the mean of their respective controls.

**Associated with Figure 3. See Tables S1 and S3 for detailed genotypes and statistical analyses.**

### Supplemental Figure #3

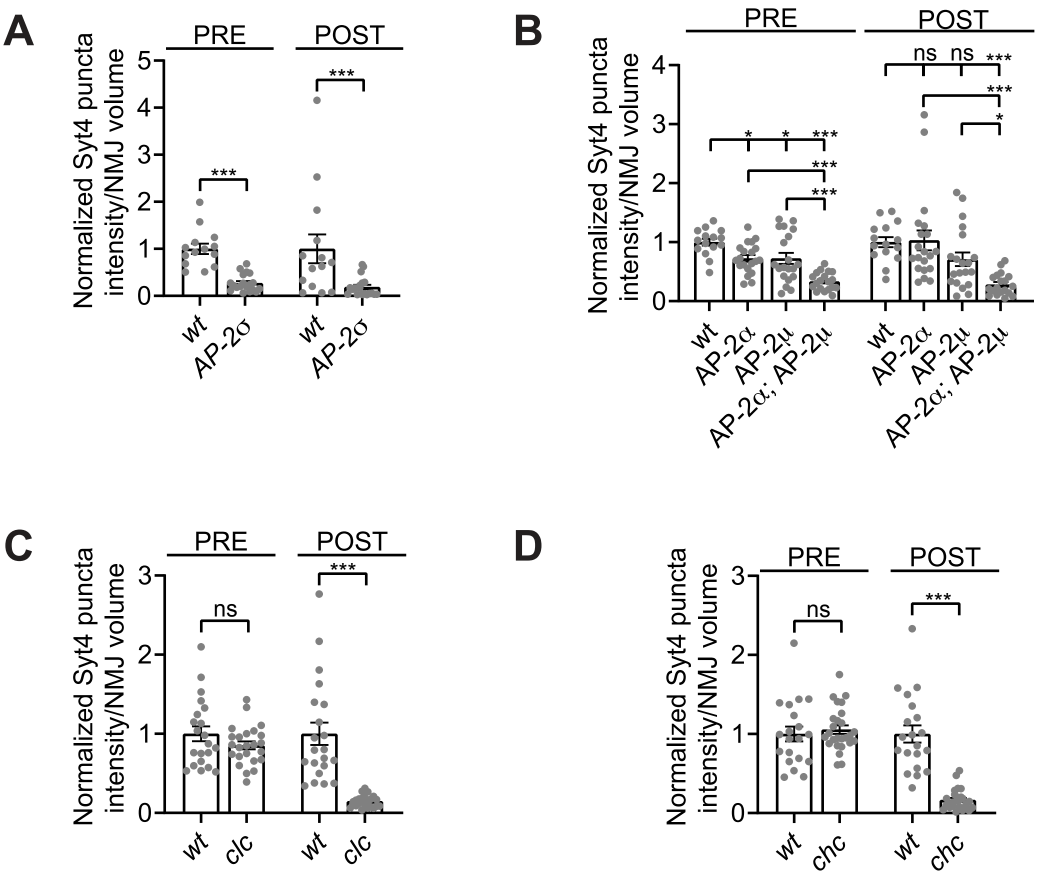


**Figure S3: EV traffic is clathrin- and AP-2-dependent.**

Fig 4A,B datasets shown with corresponding controls. (A) Quantification of Syt4-GFP pre- and postsynaptic puncta intensity in control and *AP-2σ* mutants. (B) Quantification of Syt4-GFP pre- and postsynaptic puncta intensity in control, *AP-2α*, *AP-2μ*, and *AP-2α*;*AP-2μ* mutants. (C) Quantification of Syt4-GFP pre- and postsynaptic puncta intensity in control and *clc* mutants. (D) Quantification of Syt4-GFP pre- and postsynaptic puncta intensity in control and *chc* mutants.

Data is represented as mean +/- s.e.m.; n represents NMJs. NMJ intensity measurements were normalized to presynaptic volume; all measurements were further normalized to the mean of their respective controls. **Associated with Figure 4. See Tables S1 and S3 for detailed genotypes and statistical analyses.**

**Table S1: Fly Strains**

Bloomington Drosophila Stock Center stock numbers denoted with BL; chromosome for transgene insertion indicated in roman numerals.

| **Experimental Models: Organism/allele** | | |
|---|---|---|
| GAL4$^{C155}$ (X) | (Lin and Goodman, 1994) | Flybase ID: FBti0002575 BL548 |
| GAL4$^{C380}$ (X) | (Budnik et al., 1996) | Flybase ID: FBti0016294 |
| GAL4$^{C57}$ (III) | (Budnik et al., 1996) | Flybase ID: FBti0016293 |
| GAL4$^{Vglut}$ (X) | (Daniels et al., 2008) | Flybase ID: FBti0129146 |
| *w$^{1118}$* | (Hazelrigg et al., 1984) | Flybase ID: FBal0018186 |
| *nwk$^{1}$* | (Coyle et al., 2004) | Flybase ID: FBal0154819 |
| *nwk$^{2}$,h* | (Coyle et al., 2004) | Flybase ID: FBal0154818 |
| Syt4-EGFP-KI | (Walsh et al., 2021) | |
| UAS-Nwk-22D (II) | (Rodal et al., 2008) | Flybase ID: FBal0154814 |
| UAS-Evi-EGFP (II) | (Bartscherer et al., 2006) | Flybase ID: FBal0194740 |
| UAS-APP-EGFP (II) | (Walsh et al., 2021) | |
| UAS-Tkv-mCherry (II) | (Deshpande et al., 2016) | Flybase ID: FBal0322957 |
| UAS-APP-N-myc (II) | (Merdes et al., 2004) | Flybase ID: FBal0138483 BL33798 |
| UAS-BACE (II) | (Greeve et al., 2004) | Flybase ID: FBti0139951 BL33798 |
| *synj$^{1}$* | (Verstreken et al., 2003) | Flybase ID: FBal0155922 BL24883 |
| *synj$^{2}$* | (Verstreken et al., 2003) | Flybase ID: FBal0155920 BL24884 |
| yw; UAS-Dap160-RNAi$^{JF01918}$ (III) | (Perkins et al., 2015) | Flybase ID: FBal0219906 BL25879 |
| UAS-mCherry-RNAi (III) | (Perkins et al., 2015) | Flybase ID: FBal0260847 BL35785 |
| UAS-Dcr2 (II) | (Dietzl et al., 2007) | Flybase ID: FBal0211026 BL24650 |
| *SH3PX1$^{10A}$* | (Ukken et al., 2016) | Flybase ID: FBal0318727 |
| UAS-EndoA-RNAi$^{JF02758}$ (III) | (Perkins et al., 2015) | Flybase ID: |

| | | |
|---|---|---|
| | | FBal0242340<br>BL27679 |
| UAS-Shibire$^{K44A}$ (II and X) | (Moline et al., 1999) | Flybase ID:<br>FBal0101154<br>BL5811 |
| *AP-2α*$^{40\text{-}31}$ | (Windler and Bilder, 2010) | Flybase ID:<br>FBal0219035<br>BL64752 |
| *AP-2α*$^{06694}$ | (Gonzalez-Gaitan and Jackle, 1997) | Flybase ID:<br>FBal0008113<br>BL12319 |
| *AP-2μ*$^{G7085}$ | | Flybase ID:<br>FBal0246891<br>BL32610 |
| *AP-2μ*$^{G4842}$ | | Flybase ID:<br>FBal0241410<br>BL30102 |
| *AP-2σ*$^{KG02457}$ | (Windler and Bilder, 2010) | Flybase ID:<br>FBal0131822<br>BL13478 |
| Df(3R)ED10838 (removes *AP-2σ*) | | Flybase ID:<br>FBab0044376<br>BL9485 |
| *Clc*$^{DP00280}$ | | Flybase ID:<br>FBal0216543<br>BL21806 |
| *Chc*$^{B}$ | | Flybase ID:<br>FBal0326290<br>BL67112 |
| *Vps35*$^{e42}$ | (Port et al., 2008) | Flybase ID:<br>FBal0221801 |
| Df(2R)Exel6078 (removes *Vps35*) | | Flybase ID:<br>FBab0038016<br>BL7558 |
| UAS-DCTN1-p150Δ96B (II) | (Allen et al., 1999) | Flybase ID:<br>FBal0102986<br>BL51645 |


Allen, M.J., Shan, X., Caruccio, P., Froggett, S.J., Moffat, K.G., and Murphey, R.K. (1999). Targeted expression of truncated glued disrupts giant fiber synapse formation in *Drosophila*. J Neurosci *19*, 9374-9384.

Bartscherer, K., Pelte, N., Ingelfinger, D., and Boutros, M. (2006). Secretion of Wnt ligands requires Evi, a conserved transmembrane protein. Cell *125*, 523-533.

Budnik, V., Koh, Y.H., Guan, B., Hartmann, B., Hough, C., Woods, D., and Gorczyca, M. (1996). Regulation of synapse structure and function by the *Drosophila* tumor suppressor gene dlg. Neuron *17*, 627-640.

Coyle, I.P., Koh, Y.H., Lee, W.C., Slind, J., Fergestad, T., Littleton, J.T., and Ganetzky, B. (2004). Nervous wreck, an SH3 adaptor protein that interacts with Wsp, regulates synaptic growth in *Drosophila*. Neuron *41*, 521-534.

Daniels, R.W., Gelfand, M.V., Collins, C.A., and DiAntonio, A. (2008). Visualizing glutamatergic cell bodies and synapses in *Drosophila* larval and adult CNS. J Comp Neurol *508*, 131-152.

Deshpande, M., Feiger, Z., Shilton, A.K., Luo, C.C., Silverman, E., and Rodal, A.A. (2016). Role of BMP receptor traffic in synaptic growth defects in an ALS model. Mol Biol Cell *27*, 2898-2910.

Dietzl, G., Chen, D., Schnorrer, F., Su, K.C., Barinova, Y., Fellner, M., Gasser, B., Kinsey, K., Oppel, S., Scheiblauer, S., *et al.* (2007). A genome-wide transgenic RNAi library for conditional gene inactivation in *Drosophila*. Nature *448*, 151-156.

Gonzalez-Gaitan, M., and Jackle, H. (1997). Role of *Drosophila* alpha-adaptin in presynaptic vesicle recycling. Cell *88*, 767-776.

Hazelrigg, T., Levis, R., and Rubin, G.M. (1984). Transformation of *white* locus DNA in *Drosophila*: dosage compensation, zeste interaction, and position effects. Cell *36*, 469-481.

Lin, D.M., and Goodman, C.S. (1994). Ectopic and increased expression of Fasciclin II alters motoneuron growth cone guidance. Neuron *13*, 507-523.

Merdes, G., Soba, P., Loewer, A., Bilic, M.V., Beyreuther, K., and Paro, R. (2004). Interference of human and *Drosophila* APP and APP-like proteins with PNS development in *Drosophila*. EMBO J *23*, 4082-4095.

Moline, M.M., Southern, C., and Bejsovec, A. (1999). Directionality of wingless protein transport influences epidermal patterning in the *Drosophila* embryo. Development *126*, 4375-4384.

Parks, A.L., Cook, K.R., Belvin, M., Dompe, N.A., Fawcett, R., Huppert, K., Tan, L.R., Winter, C.G., Bogart, K.P., Deal, J.E., *et al.* (2004). Systematic generation of high-resolution deletion coverage of the *Drosophila melanogaster* genome. Nat Genet *36*, 288-292.

Perkins, L.A., Holderbaum, L., Tao, R., Hu, Y., Sopko, R., McCall, K., Yang-Zhou, D., Flockhart, I., Binari, R., Shim, H.S., *et al.* (2015). The Transgenic RNAi Project at Harvard Medical School: Resources and Validation. Genetics *201*, 843-852.

Port, F., Kuster, M., Herr, P., Furger, E., Banziger, C., Hausmann, G., and Basler, K. (2008). Wingless secretion promotes and requires retromer-dependent cycling of Wntless. Nat Cell Biol *10*, 178-185.

Rodal, A.A., Motola-Barnes, R.M., and Littleton, J.T. (2008). Nervous Wreck and Cdc42 cooperate to regulate endocytic actin assembly during synaptic growth. J Neurosci *28*, 8316-8325.

Ukken, F.P., Bruckner, J.J., Weir, K.L., Hope, S.J., Sison, S.L., Birschbach, R.M., Hicks, L., Taylor, K.L., Dent, E.W., Gonsalvez, G.B., *et al.* (2016). BAR-SH3 Sorting nexins are conserved Nervous wreck interactors that organize synapses and promote neurotransmission. J Cell Sci *129*, 166-177.

Verstreken, P., Koh, T.W., Schulze, K.L., Zhai, R.G., Hiesinger, P.R., Zhou, Y., Mehta, S.Q., Cao, Y., Roos, J., and Bellen, H.J. (2003). Synaptojanin is recruited by endophilin to promote synaptic vesicle uncoating. Neuron *40*, 733-748.

Walsh, R.B., Dresselhaus, E.C., Becalska, A.N., Zunitch, M.J., Blanchette, C.R., Scalera, A.L., Lemos, T., Lee, S.M., Apiki, J., Wang, S., *et al.* (2021). Opposing functions for retromer and Rab11 in extracellular vesicle traffic at presynaptic terminals. J Cell Biol *220*.

Windler, S.L., and Bilder, D. (2010). Endocytic internalization routes required for delta/notch signaling. Curr Biol *20*, 538-543.

#### Table S2: Antibodies

| REAGENT | SOURCE | IDENTIFIER | Concentration |
|---|---|---|---|
| **Antibodies** | | | |
| α-HRP-488, -RRX, -647 | Jackson ImmunoResearch | | 1:250-1:500 (IHC) |
| α–GFP nanobody | Nanotag Biotechnologies | N0304 | 1:250 (IHC) |
| α-Synaptotagmin-1 | (West et al., 2015) | | 1:1000 (IHC) |
| α-Dlg | (Parnas et al., 2001), DSHB | 4F3 | (IHC) |
| α-Nrg | (Hortsch et al., 1990), DHSB | BP104 | 1:100 (IHC) |
| α-APP-CTFs | Sigma | A8717 | 1:5000 (WB) |
| α-actin | (Lin, 1981), DHSB | JLA20 | 1:100 (WB) |
| α-even-skipped | (Patel, Condron & Zinn, 1994) DHSB | 2B8 | 1:30 (IHC) |

Hortsch, M., Bieber, A.J., Patel, N.H., and Goodman, C.S. (1990). Differential splicing generates a nervous system-specific form of *Drosophila* neuroglian. Neuron *4*, 697-709.
Lin, J.J. (1981). Monoclonal antibodies against myofibrillar components of rat skeletal muscle decorate the intermediate filaments of cultured cells. Proc Natl Acad Sci U S A *78*, 2335-2339.
Parnas, D., Haghighi, A.P., Fetter, R.D., Kim, S.W., and Goodman, C.S. (2001). Regulation of postsynaptic structure and protein localization by the Rho-type guanine nucleotide exchange factor dPix. Neuron *32*, 415-424.
West, R.J., Lu, Y., Marie, B., Gao, F.B., and Sweeney, S.T. (2015). Rab8, POSH, and TAK1 regulate synaptic growth in a *Drosophila* model of frontotemporal dementia. J Cell Biol *208*, 931-947.

#### Table S3: Statistics by Dataset

Experiments were done at m67 unless otherwise noted
Presynaptic volume: α-HRP objects > 7μm$^3$
Postsynaptic volume: 3μm dilated from presynaptic volume
A: Sum intensity of signal in thresholded objects in presynaptic volume, normalized to presynaptic volume
B: Sum intensity of signal in thresholded objects in postsynaptic volume, normalized to presynaptic volume

| Figure | Genotype/Conditions | N | Measurement | Statistical Test(s) |
|---|---|---|---|---|
| 1A | $GAL4^{C380}$ / Y ; Syt4-EGFP<br><br>$GAL4^{C380}$ / Y ; ; *h, nwk*$^2$ Syt4-EGFP | 22 NMJs from 6 animals<br><br>18 NMJs from 6 animals | B | Mann-Whitney ($p<0.0001$) |
| 1B | $GAL4^{C380}$ / Y ; Syt4-EGFP<br><br>$GAL4^{C380}$ / Y ; ; *h, nwk*$^2$ Syt4-EGFP<br><br>$GAL4^{C380}$ / Y ; UAS-$Nwk^{22D}$ / + ; *h, nwk*$^2$, Syt4-EGFP | 23 NMJs from 6 animals<br><br>20 NMJs from 6 animals<br><br>21 NMJs from 6 animals | A<br><br>B<br><br>Number of postsynaptic puncta, normalized to presynaptic volume | Pre: Kruskal-Wallis with Dunn's multiple comparisons<br><br>($p<0.0001$)<br><br>Post: One-way ANOVA with Tukey's multiple comparisons<br><br>($p<0.0001$)<br><br>Puncta number: One-way ANOVA with Tukey's multiple comparisons<br><br>($p<0.0001$) |
| 1C | $GAL4^{C155}$; UAS-Evi-long-GFP / +<br><br>$GAL4^{C155}$; UAS-Evi-long-GFP / + ; *nwk*$^1$ / *h, nwk*$^2$ | 24 NMJs from 8 animals<br><br>22 NMJs from 8 animals | A<br><br>B<br><br>Number of postsynaptic puncta, normalized to presynaptic volume | (both measurements)<br><br>Pre: Mann-Whitney<br><br>($p<0.0001$) |

| | | | | |
|---|---|---|---|---|
| | | | | Post: Mann-Whitney<br><br>($p<0.0001$) |
| 1C | Gal4$^{C155}$ / Y ; UAS-hAPP-EGFP / +<br><br>Gal4$^{C155}$ / Y ; UAS-hAPP-EGFP / + ; *nwk*$^{1}$ / *h, nwk*$^{2}$ | 14 NMJs from 6 animals<br><br>13 NMJs from 5 animals | A<br><br>B | Pre: Mann-Whitney<br><br>($p=0.0004$)<br><br>Post: Mann-Whitney<br><br>($p<0.0001$) |
| 1C | *w*$^{1118}$<br><br>*nwk*$^{1}$ / *h, nwk*$^{2}$ | 14 NMJs from 5 animals<br><br>10 NMJs from 4 animals | A<br><br>B | Pre: t-test<br><br>($p<0.0001$)<br><br>Post: t-test<br><br>($p=0.0003$) |
| 1C | Syt4-EGFP<br><br>*h, nwk*$^{2}$, Syt4-EGFP | 22 NMJs from 8 animals<br><br>16 NMJs from 7 animals | A<br><br>B | Pre: t-test<br><br>($p<0.0001$)<br><br>Post: Mann-Whitney<br><br>($p<0.0001$) |
| 1C | Gal4$^{C155}$ / Y ; UAS-hAPP-EGFP / +<br><br>Gal4$^{C155}$ / Y ; UAS-hAPP-EGFP / + ; *nwk*$^{1}$ / *h, nwk*$^{2}$ | 17 NMJs from 8 animals<br><br>19 NMJs from 8 animals | A<br><br>B | Pre: Mann-Whitney<br><br>($p<0.0001$)<br><br>Post: Mann-Whitney<br><br>($p<0.0001$) |
| 1D | Gal4$^{C155}$ / Y or Gal4$^{C155}$ / w1118 | 22 NMJs<br><br>22 NMJs | A<br><br>B | t-test<br><br>($p=0.7528$) |

| | | | | |
|---|---|---|---|---|
| | Gal4$^{C155}$ / Y or Gal4$^{C155}$ / *w*$^{1118}$; *nwk*$^{1}$ / *h, nwk*$^{2}$ | | | |
| 1E | Gal4$^{vglut}$ / Y ; UAS-Tkv-mCherry / +<br><br>Gal4$^{vglut}$ / Y ; UAS-Tkv-mCherry / + ; *h, nwk*$^{2}$ | 15 NMJs from 7 animals<br><br>18 NMJs from 7 animals | A<br><br>B | t-test<br><br>(p=0.6963) |
| 1F | Syt4-EGFP<br><br>*nwk*$^{2}$, Syt4-EGFP | 5 brains<br><br>7 brains | Mean intensity of VG signal per VG volume | Mann-Whitney<br><br>(p=0.3434) |
| 1F | Syt4-EGFP<br><br>*nwk*$^{1}$, Syt4-EGFP | 8 brains<br><br>8 brains | Mean intensity of neuropil signal per neuropil area | t-test<br><br>(p=0.3618) |
| 1F | GAL4$^{Vglut}$ / Y; UAS-hAPP-EGFP / +<br><br>GAL4$^{Vglut}$ / Y; UAS-hAPP-EGFP / +; *nwk*$^{1}$ / *h, nwk*$^{2}$ | 17 NMJs from 8 animals<br><br>18 NMJs from 8 animals | Sum intensity of axon signal per axon area | Mann-Whitney<br><br>(p = 0.2066) |
| 2AB | w$^{1118}$<br><br>*nwk*$^{1}$ / *h, nwk*$^{2}$<br><br>GAL4$^{C57}$ / + ; UAS-Nwk-22D / + ; *nwk*$^{1}$ / *h, nwk*$^{2}$<br><br>GAL4$^{Vglut}$ / + ; UAS-Nwk-22D / + ; *nwk*$^{1}$ / *h, nwk*$^{2}$ | 14 NMJs<br><br>13 NMJs<br><br>12 NMJs<br><br>12 NMJs | Fold change in mEJP frequency and fold change in mEJP frequency 100s after stimulus | One-way ANOVA with Tukey's multiple comparisons<br><br>(p<0.0001) |
| 2CD | *w*$^{1118}$<br><br>*nwk*$^{1}$ / *h, nwk*$^{2}$ | 15 NMJs<br><br>15 NMJs | Number of ghost boutons | wt: t-test<br><br>(p<0.0001)<br><br>nwk: t-test<br><br>(p = 0.1295) |

| | | | | |
|---|---|---|---|---|
| 2E | Gal4$^{C155}$ / Y and Gal4$^{C155}$ / *w*$^{1118}$<br><br>*w*$^{1118}$/ Y and + ; UAS-hAPP-695-N-myc, UAS BACE1 / +<br><br>Gal4$^{C155}$ / Y and + ; UAS-hAPP-695-N-myc, UAS-BACE1 / +<br><br>Gal4$^{C155}$ / Y and + ; *nwk*$^{1}$ / *h, nwk*$^{2}$<br><br>Gal4$^{C155}$ / Y and + ; UAS-hAPP-695-N-myc, UAS-BACE1 / + ; *h, nwk*$^{2}$<br><br>Gal4$^{C155}$ / Y and + ; UAS-hAPP-695-N-myc, UAS-BACE1 / + ; *h, nwk*$^{2}$ / + | 1657 pupal cases<br><br>1509 pupal cases<br><br>2479 pupal cases<br><br>1736 pupal cases<br><br>1477 pupal cases<br><br>1989 pupal cases | Total N eclosed over total pupal cases | Chi-Squared<br><br>($p<0.0001$) |
| 2F | Gal4$^{C155}$ / Y<br><br>Gal4$^{C155}$ / Y ; UAS-hAPP-695-N-myc / +<br><br>Gal4$^{C155}$ / Y ; UAS-hAPP-695-N-myc / + ; *h, nwk*$^{2}$<br><br>Gal4$^{C155}$ / Y ; UAS-hAPP-695-N-myc, UAS-BACE1 / +<br><br>Gal4$^{C155}$ / Y ; UAS-hAPP-695-N-myc, UAS-BACE1 / + ; *h, nwk*$^{2}$ | 3 replicates of 15 fly heads | APP CTF intensity normalized to JLA20 (actin) intensity | One-way ANOVA with Tukey's multiple comparisons<br><br>($p=0.1336$) |
| 3A | Syt4-EGFP<br><br>*synj*$^{1}$/*synj*$^{2}$ ; Syt4-EGFP | 24 NMJs from 7 animals<br><br>19 NMJs from 6 animals | A<br><br>B | Pre: t-test<br><br>($p<0.0001$)<br><br>Post: t-test<br><br>($P<0.0001$) |

| | | | | |
|---|---|---|---|---|
| 3A | GAL4$^{C380}$ / Y ; UAS-dicer / + ; UAS-mCherry RNAi / Syt4-EGFP<br><br>GAL4$^{C380}$ / Y ; UAS-dicer / + ; UAS-Dap160 RNAi / Syt4-EGFP | 19 NMJs from 6 animals<br><br>16 NMJs from 6 animals | A<br><br>B | Pre: t-test<br><br>($p=0.0066$)<br><br>Post: Mann-Whitney<br><br>($p<0.0001$) |
| 3A | Syt4-EGFP<br><br>*SH3PX1*$^{10A}$ , Syt4-EGFP | 23 NMJs from 7 animals<br><br>20 NMJs from 7 animals | A<br><br>B | Pre: t-test<br><br>($p<0.0001$)<br><br>Post: Mann-Whitney<br><br>($p<0.0001$) |
| 3B | GAL4$^{C155}$ / Y<br><br>GAL4$^{C155}$ / Y ; UAS-*Shi*$^{K44A}$ / + | 12 NMJs from 6 animals<br><br>14 NMJs from 6 animals | B | Mann-Whitney<br><br>($p<0.0001$) |
| 3B | GAL4$^{C380}$ / + ; Syt4-EGFP<br><br>GAL4$^{C380}$ / + ; UAS-endoA RNAi Syt4-EGFP / SYT4 EGFP | 11 NMJs from 3 animals<br><br>20 NMJs from 7 animals | B | Mann-Whitney<br><br>($p=0.0011$) |
| 4A | Syt4-EGFP / +<br><br>*AP2σ*$^{KG02457}$, Syt4-EGFP / Df(ED10838) | 14 NMJs from 6 animals<br><br>19 NMJs from 5 animals | A<br><br>B | Pre: t-test<br><br>($p<0.0001$)<br><br>Post: Mann-Whitney<br><br>($p=0.0006$) |
| 4A | Syt4-EGFP | 15 NMJs from 6 animals | A<br><br>B | Pre: One-way ANOVA with Tukey’s multiple comparison |

| | | | | |
|---|---|---|---|---|
| | *AP-2α*$^{40-31}$/ *AP-2α*$^{06694}$ ; Syt4-EGFP<br><br>*AP-2μ*$^{G7085}$ Syt4-EGFP / *AP-2μ*$^{G4842}$ Syt4-EGFP<br><br>*AP-2α*$^{40-31}$/ *AP-2α*$^{06694}$ ; *AP-2μ*$^{G7085}$ Syt4-EGFP / *AP-2μ*$^{G4842}$ Syt4-EGFP | 20 NMJs from 7 animals<br><br>20 NMJs from 6 animals<br><br>19 NMJs from 6 animals | | ($p<0.0001$)<br><br>Post: Kruskal-Wallis<br><br>($p<0.0001$) |
| 4BC | Syt4-EGFP<br><br>*clc*$^{DP00280}$ , Syt4 EGFP<br><br>20°C | 21 NMJs from 7 animals<br><br>24 NMJs from 7 animals | A<br><br>B | Pre: t-test<br><br>($p=0.1633$)<br><br>Post: Mann Whitney<br><br>($p<0.0001$) |
| 4B | Syt4-EGFP<br><br>*chc*$^{B}$, Syt4-EGFP<br><br>20°C | 20 NMJs from 8 animals<br><br>26 NMJs from 8 animals | A<br><br>B | Pre: t-test<br><br>($p=0.5870$)<br><br>Post: Mann Whitney<br><br>($p<0.0001$) |
| 4D | Syt4-EGFP<br><br>*h, nwk*$^{2}$ *,* Syt4-EGFP<br><br>*clc*$^{DP00280}$ Syt4-EGFP | 16 NMJs from 6 animals<br><br>19 NMJs from 6 animals<br><br>16 NMJs from 6 animals<br><br>17 NMJs from 6 animals | A<br><br>B | Pre: One-way ANOVA with Tukey's multiple comparisons<br><br>($p<0.0001$)<br><br>Post: Kruskal-Wallis with Dunn's multiple comparisons test<br><br>($p<0.0001$) |

| | | | | |
|---|---|---|---|---|
| | *h, nwk*$^{2}$, *clc*$^{DP00280}$ Syt4-EGFP<br><br>20°C | | | |
| 5B | Syt4-EGFP<br><br>*vps35*$^{e42}$/Df6078; Syt4-EGP<br><br>*h, nwk*$^{2}$ *,* Syt4-EGFP<br><br>vps35[e42]/Df6078; *h, nwk*$^{2}$, Syt4-EGFP | 21 NMJs from 6 animals<br><br>20 NMJs from 6 animals<br><br>19 NMJs from 6 animals<br><br>22 NMJs from 6 animals | A<br><br>B<br><br>Number of postsynaptic Syt4 puncta normalized to presynaptic volume<br><br>Mean volume of postsynaptic Syt4 puncta | Pre: Kruskal-Wallis with Dunn's multiple comparisons<br><br>(p<0.0001)<br><br>Post: One-way ANOVA with Tukey's multiple comparisons<br><br>(p<0.0001)<br><br>Puncta number: One-way ANOVA with Tukey's multiple comparisons<br><br>(p<0.0001)<br><br>Puncta volume: One-way ANOVA with Tukey's multiple comparisons<br><br>(p<0.0001) |
| 5C | GAL4$^{C380}$ / + ; Syt4-EGFP<br><br>GAL4$^{C380}$ / + ; UAS-DCTN$^{p150}$ / + ; Syt4-EGFP<br><br>GAL4$^{C380}$ / +; *h, nwk*$^{2}$ Syt4-EGFP<br><br>GAL4$^{C380}$ / + ; UAS-DCTN$^{p150}$ / + ; *h, nwk*$^{2}$ Syt4-EGFP | 19 NMJs from 6 animals<br><br>17 NMJs from 6 animals<br><br>17 NMJs from 6 animals<br><br>18 NMJs from 6 animals | A<br><br>B | Pre: Kruskal-Wallis with Dunn's multiple comparisons<br><br>(p<0.0001)<br><br>Post: One-way ANOVA with Tukey's multiple comparisons<br><br>(p<0.0001) |
